# Accurate prediction of protein torsion angles using evolutionary signatures and recurrent neural network

**DOI:** 10.1101/2021.05.06.442265

**Authors:** Yong-Chang Xu, Tian-Jun ShangGuan, Xue-Ming Ding, Ngaam J. Cheung

## Abstract

The amino acid sequence of a protein contains all the necessary information to specify its shape, which dictates its biological activities. However, it is challenging and expensive to experimentally determine the three-dimensional structure of proteins. The backbone torsion angles, as an important structural constraint, play a critical role in protein structure prediction, and accurately predicting the angles can considerably advance the tertiary structure prediction by accelerating efficient sampling of the large conformational space for low energy structures. On account of the rapid growth of protein databases and striking breakthroughs in deep learning algorithms, computational advances allow us to extract knowledge from large-scale data to address key biological questions. Here we propose evolutionary signatures that are computed from protein sequence profiles, and a deep neural network, termed ESIDEN, that adopts a straightforward architecture of recurrent neural networks with a small number of learnable parameters. The proposed ESIDEN is validated on three benchmark datasets, including D2020, TEST2016/2018, and CASPs datasets. On the D2020, using the combination of the four novel features and basic features, the ESIDEN achieves the mean absolute error (MAE) of 15.7 and 19.8 for *ϕ* and *ψ*, respectively. Comparing to the best-so-far methods, we show that the ESIDEN significantly improves the angle *ψ* by the MAE decrements of more than 3.5 degrees on both TEST2016 and TEST2018 and achieves better MAE of the angle *ϕ* by decrements of at least 0.3 degrees although it adopts simple architecture and fewer learnable parameters. On fifty-nine template-free modeling targets, the ESIDEN achieves high accuracy by reducing the MAEs by 0.6 and more than 2.3 degrees on average for the torsion angles *ϕ* and *ψ* in the CASPs, respectively. Using the predicted torsion angles, we infer the tertiary structures of four representative template-free modeling targets that achieve high precision with regard to the root-mean-square deviation and TM-score by comparing them to the native structures. The results demonstrate that the ESIDEN can make accurate predictions of the torsion angles by leveraging the evolutionary signatures. The proposed evolutionary signatures would be also used as alternative features in predicting residue-residue distance, protein structure, and protein-ligand binding sites. Moreover, the high-precision torsion angles predicted by the ESIDEN can be used to accurately infer protein tertiary structures, and the ESIDEN would potentially pave the way to improve protein structure prediction.

## 1 Introduction

Proteins play important roles in biological activities, and their functional significance is determined by their three-dimensional structure. However, it is difficult and expensive to experimentally determine protein tertiary structures. Moreover, with the rapid large-scale sequencing technologies, a gap between the huge number of protein sequences and a small number of known structures is being enlarged. Predicting protein three-dimensional structures is an alternative way to narrow the gap. It has been a grand challenge to make an accurate prediction without any structural information in computational biophysics for decades^1, 2^, as there are mainly two difficulties in the prediction: (1) efficient sampling methods to search an astronomically larger conformation space^3^, and (2) accurate free energy determination to find the most stable shape^1^. Although it is extremely hard to search the large space of possible structures for the one with the lowest energy, inferred structural constraints, such as contact/distance between pairwise residues, have advanced protein structure prediction and decreased the deviation between the predicted and the authentic structures^4–6^. The backbone torsion angles, as an important structural constraint, also play a critical role in protein structure prediction (e.g., sampling the space of the torsion angles (*ϕ, ψ*) to investigate protein folding^7^) and refinement^8^, and they are also commonly used as constraints in many computational methods, e.g., CNS^9^, CYANA^10^, and AMBER^11^ to determine protein structures. Accurately predicting the torsion angles can considerably advance the tertiary structure prediction by accelerating efficient sampling of the large conformational space for the low energy structures.

Owing to the larger protein databases and the development of computing resources, as well as advances in machine learning methods and deep neural networks, the accuracy of protein backbone torsion angle prediction has been improved increasingly. Typically, machine learning-based methods including neural networks^12–14^, support vector machines (SVM)^13–15^, and hidden Markov models^16,17^ predict discrete states of *ϕ /ψ* angle values. Recently, computational advances have been developed to predict real values of the torsion angles. The DESTRUCT method uses position specific scoring matrix (PSSM) to build iterative neural network models for the first time predicting the real value of the angle *ψ* although the correlation coefficient between the predicted and the real value is less than 0.5^18^. The Real-SPINE model was designed based on integrated neural networks to improve the correlation coefficient for the angle *ψ* to more than 0.6^12^. Based on a composite machine-learning algorithm, Wu et al. developed the ANGLOR^13^ to separately predict *ϕ* using the feed-forward neural network and *ψ* using the SVM.

Considerable progress has recently been made by leveraging computational advances, especially deep learning (DL), in both protein secondary and tertiary structure prediction. For example, DL-based approaches have been convincingly demonstrated to predict structural constraints that successfully guide protein folding^5,6^. The SPIDER2 method was developed to predict the torsion angles using iterative neural networks^19^, while the SPIDER3^20^ takes advantage of the long short-term memory bidirectional recurrent neural networks (LSTM-BRNN^21^) to remove the effect of the sliding window that was used in the SPIDER2^19^. The DeepRIN^22^ was designed based on the architectures of the Inception^23^ and the ResNet^24^ networks. Gao et al. developed a deep neural network-based model to predict discrete angles within 5° bins^25^. As a hybrid method, the RaptorX-Angle combines K-means clustering and deep learning techniques to predict real-valued angles^26^, as claimed it takes advantage of both discrete and continuous representation of the torsion angles. The SPOT-1D^27^ is a hybrid model that employs an ensemble of LSTM-BRNN and ResNet. The OPUS-TASS was developed based on the network architecture of the modified transformer and CNN modules, and it is trained using the additional feature (PSP19)^8^. The same network in the OPUS-TASS was trained for six different tasks including secondary structure, backbone torsion angles (TA), discrete descriptors of local backbone structure (CSF3), solvent accessible surface area (ASA), and side-chain dihedral angles (SDA).

Advances in predicting protein torsion angles have also benefited increasingly from machine learning-based methods. As a growing focus on DL-based methods, accurate prediction of the torsion angle is not only dependent on architecture of DL but also on information (features) extracted from protein sequences. In most cases, the performances of a DL-based method are highly determined by the information. Generally, classical features, such as PSSM^28^, physicochemical properties (PP^29^), and amino acid (AA), have been widely and successfully used to predict protein secondary/tertiary structure, but the classical features are far from satisfactory for existing DL-based models as a large number of parameters needs to be learned for better predictions. Moreover, the deeper the DL model is, the more difficult we optimize its parameters. To meet these requirements, we firstly present four evolutionary signatures as novel features, including the relative entropy (RE), the degree of conservation (DC), the position-specific substitution probabilities (PSSP), and the Ramachandran basin potential (RBP) statistically derived from protein sequences by removing redundant or unnecessary signals, and we also develop an evolutionary signatures-driven deep neural network (termed ESIDEN) that adopts straightforward architecture of recurrent neural networks to improve the prediction accuracy of the torsion angles.

## 2 Materials and methods

In this section, we describe the benchmark datasets and the novel features used in our study, and the proposed method is presented in detail.

### 2.1 Datasets and input features

Two benchmark datasets are used for both training and test in our study, and they include a culled dataset (D2020, Table S1) from the PISCES^30^ and the SPOT-1D dataset^27,31^ (Table S2). The D2020 dataset was culled from the PISCES server^30^ with less than 25% identity and less than 1.6Å resolution (R-factor is 0.25) (released in December 2020), which contains 8,669 protein chains. We filter out the protein whose sequence length less than 500 and finally obtain 7,443 proteins (Table S1), and these proteins are randomly classified into three groups by percentage 0.8:0.1:0.1, that is, 5,995 proteins in the training dataset, 744 proteins in the validation dataset, and the rest 744 proteins for the test dataset. For each protein chain, its torsion angles (*ϕ* and *ψ*) are extracted by using the *stride*^32^ from its structure file that is downloaded from the Protein Data Bank (PDB)^33^. The angles in the training dataset are used to train the proposed network, while the validation and test datasets are used to evaluate and measure the performance of the built model. On the other hand, we use the SPOT-1D dataset^27,31^ (Table S2) for comparing the ESIDEN to other best-so-far methods. Briefly, the dataset contains 12,450 protein chains culled from the PISCES server^30^ with a high resolution of less than 2.5Å, R-factor less than 1, and the cutoff of the sequence identity is set to 25%. By removing proteins of more than 700 amino acids, there are 10,029, 983, and 1,213 protein chains in training, validation, and test (TEST2016) datasets, respectively. An additional test dataset (TEST2018) that includes 250 protein chains is also used for fair comparison among different methods, and each protein is of the resolution less than 2.5 and R-free no more than 0.25. To further evaluate different methods, we collect 59 proteins from the template-free modeling (TFM) targets in the Critical Assessment of protein Structure Prediction (CASP). There are twenty-seven, eleven, thirteen, and eight proteins from the CASP11, CASP12, CASP13, and CASP14, respectively.

For all datasets, the features of each protein are extracted from its sequence and corresponding multiple sequence alignment (MSA). The basic features include: 20 types of amino acids (AA), seven physicochemical properties (PP)^29^, including steric parameter, polarizability, normalized van der Waals volume, hydrophobicity, isoelectric point, helix probability and sheet probability, and PSSM^28^ that is commonly used for protein secondary structure prediction^28^, residue contact prediction^31^ and torsion angles prediction^13,19,34^. The PSSM of each protein is derived from its MSA by searching its sequence against NCBI non-redundant dataset using PSI-BLAST^35^ with default parameters (e-value 0.001 and 3 iterations). In the PSSM, each amino acid has a vector that composes of 20-dimension scores, accordingly, the dimension of each PSSM is *L* × 20, where *L* is the number of amino acids of a given sequence.

The four novel features proposed in our study include the degree of conservation (DC), the relative entropy (RE), the position-specific substitution probabilities (PSSP), and the Ramachandran basin potential (RBP). To obtain the three features (DC, RE and PSSP), we firstly prepare a MSA for each protein by searching its sequence (query) against the Uniclust30 database (as of 2/2020)^36^ by HHblits^37^, and the MSA is trimmed using Leri *sequence_trim* tool^38^. Relying on the filtered MSA, we compute the DC, RE, and PSSP of each sequence as input features to train the ESIDEN. Second, the RBP is derived based on the query sequence from the potential of torsion angles (*ϕ* and *ψ*) using *Leri*^38^. The details of the four new features are presented in the following paragraphs.

#### The relative entropy (RE)

is to measure how a probability distribution of an amino acid in the MSA is different from that of another amino acid. The RE of an amino acid at the *i*th position is defined as follows,

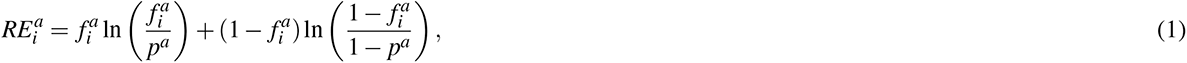

where 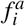 is a probability of an amino acid *a* at the *i*th position in the MSA, and *p^a^* is the background probability of the amino acid a. The dimension of each RE is *L* × 20.

#### The degree of conservation (DC)

is derived from the same MSA as that of the RE. The DC of a given amino acid *a* at the *i*th position in the MSA is defined as

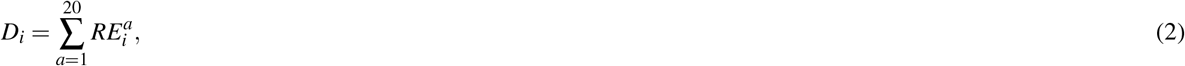

*D_i_*, of the amino acid *a* is to measure how much conservation at the *i*th position in the MSA, and, generally, it provides rich evolutionary information (e.g., conservation) of structured regions (e.g., *α*-helix and *β*-strand)^39^. The dimension of the DC is *L* × 1.

#### The position-specific substitution probabilities (PSSP)

is derived from protein sequence profiles using the evolutionary statistical energy (a Markov Random field or a Potts model in statistical physics^40,41^) that is defined as follows,

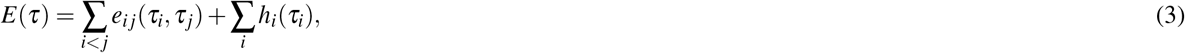

where *h_i_* and *e_ij_* are site-specific bias terms of a single amino acid and coupling terms between pairwise amino acids, respectively. Without considering inter-dependencies between pairwise amino acids, the site-specific amino acid constraints *h_i_* are used to construct the PSSP. In our study, we use the same MSA as that of RE and DC to optimize the Markov Random field by *Leri*^38^ and obtain the PSSP from the optimized site-specific bias *h_i_* (Eq. (3)). Without the gaps, the dimension of the PSSP of each protein is *L* × 20.

#### The Ramachandran basin potential (RBP)

Neighboring amino acids have been shown to exert a strong influence on protein structure^42,43^. In this study, we report for the first time that a statistical potential derived from the Ramachandran basins is used to predict the torsion angles. Briefly, the potential of torsion angles is computed from proteins with 25% sequence identity, and the potential is divided into 72 × 72 bins (5° × 5° of each bin). To generate the RBP of a given sequence, the probability distribution of each amino acid is computed by: (1) taking advantage of two close neighbors of a central residue, that is, the left and right neighbors of the triple residues, and (2) taking predicted secondary structure of the three residues into account, e.g., the Q3 (helix/sheet/coil classes, H/E/C) or Q8 (3-turn helix/4-turn helix/5-turn helix/hydrogen-bonded turn/extended strand in parallel and/or anti-parallel *β*-sheet conformation/residue in isolated *β*-bridge/bend/coil, G/H/I/T/E/B/S/C). In this study, we treat the secondary structure of each residue as A, that is, its secondary structure can be any one of Q3.

The RBP is employed as a new feature to predict protein torsion angles. As shown in Figure 1, each amino acid has its own Ramanchnadran basin that is derived from the statistical potential of the torsion angles (*ϕ* and *ψ*). It is generated by the *Leri* software, and we leverage the nonlinear dimensionality reduction technique, the t-SNE algorithm^44^, to reduce the original dimension 72 × 72 to 72 × 2 for efficient computation. In practice, the reduced RBP is flattened to a one-dimensional vector 1 × 144, and the dimension of the RBP feature is *L* × 144 of a protein chain.

**Figure 1.**
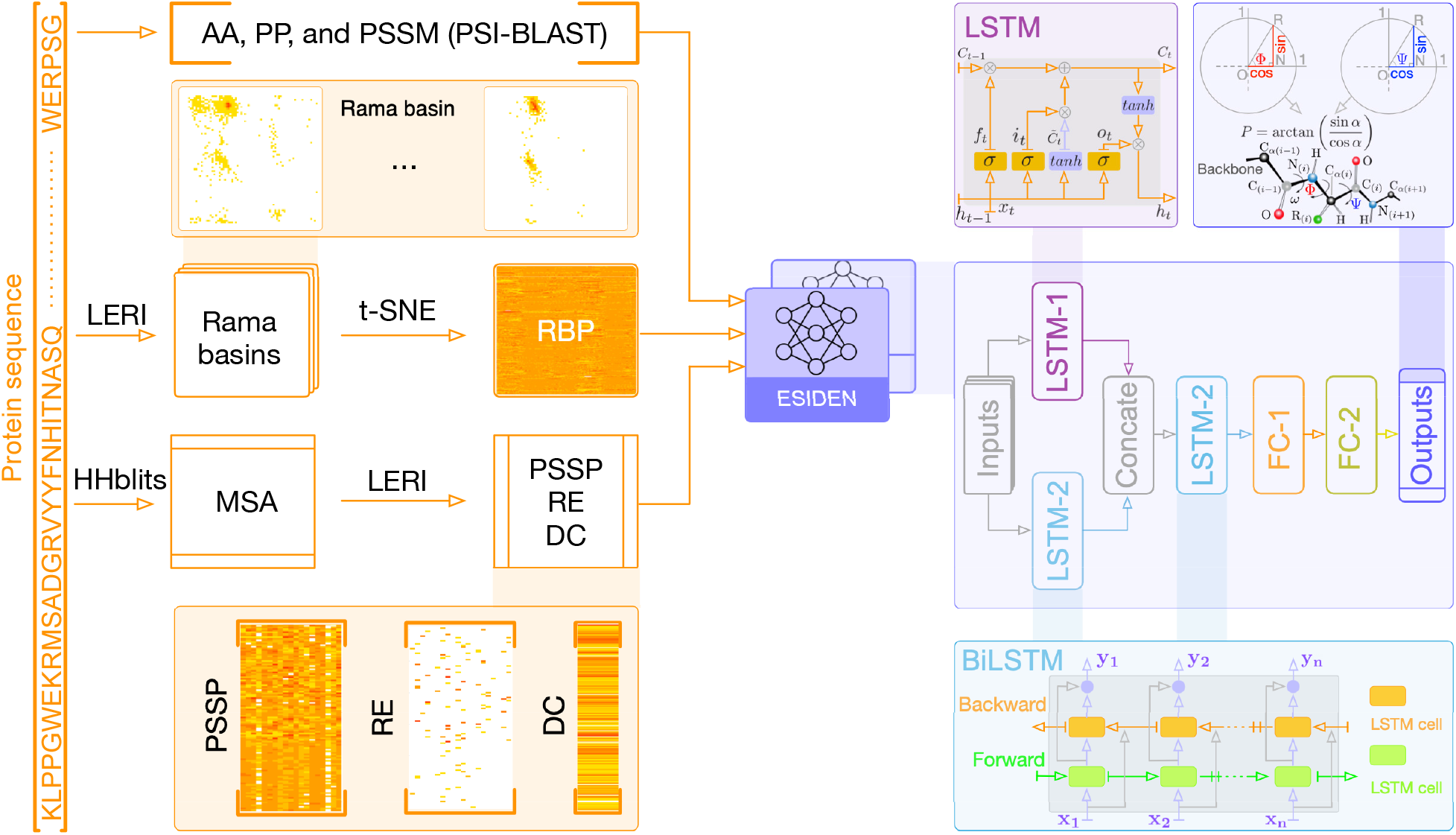
Schematics of prediction system and the ESIDEN network for protein torsion angles. Starting from the primary sequence, the system extracts and processes the features (*L* × 232) as inputs to feed the ESIDEN network. The ESIDEN mainly composes of LSTM (LSTM-1 is in parallel with LSTM-2 and they are concatenated to another LSTM-2) and FC modules.

### 2.2 The proposed system

The prediction system (Figure 1) is to estimate the torsion angles (*ϕ* and *ψ*) from protein primary sequence, and it mainly consists of two parts: (1) features extraction and processing from the primary sequence; and (2) predicting the torsion angle using the proposed method (ESIDEN). In addition to the basic features, the four novel features extracted from protein sequences are also leveraged in our network for highly accurate torsion angle prediction. There are seven features, including AA, PP, PSSM, RE, DC, PSSP, and RBP, that are derived from the primary sequence without any structural information (Figure 1). The classic features (AA, PP, and PSSM) are generated and processed into a matrix data of *L* × 47 (as discussed above), and four novel evolutionary signatures, the RE, DC, and PSSP are computed from the same MSA of each protein and processed using in-house scripts. In this study, both the new and the classic features are normalized into the range [-1,1] to enlarge differences and highlight important components of each feature. Accordingly, we achieve a feature matrix of *L* × 232 for a protein chain of *L*, and these features are fed to the proposed network (ESIDEN) to optimize its learnable parameters.

The ESIDEN is developed based on the LSTM and fully connected (FC) modules. As illustrated in Figure 1, we design the ESIDEN using the LSTM-1 and LSTM-2 in parallel, and they are concatenated to another LSTM-2 architecture, which can effectively understand the combination of different features of each protein sequence. As a basic LSTM network, each hidden layer cell in the LSTM-1 has an input that depends on the cell at the previous state. This explicit arrangement of the cell makes LSTM capture previous context information hidden in the feature of an input sequence, and it can also efficiently learning important components in the features. In the LSTM-1 module, the final output *h_t_* at the step *t* of the cell depends on other variables, including the input value 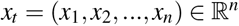, the forget gate 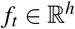, the input gate 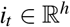, the cell state 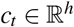, the temporary cell state 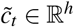 and the output gate 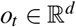 at current state, and the cell state 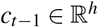 and the hidden values 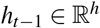 at the previous state, where the superscripts *n* and *h* are the numbers of input features and hidden units, respectively. The relationships among those variables are formulated as follows,

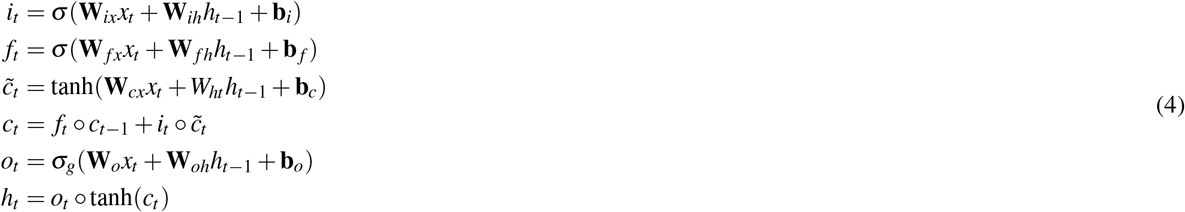

where ***σ***(·) is the Sigmoid function, and tanh is a nonlinear transformation function. **W** and **b** represent the weight matrix and bias vector of different activation units (e.g. 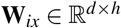 represent the input gate weight matrix), respectively. The operator o denotes the Hadamard product.

In the ESIDEN, the LSTM-2 adopts an architecture of BiLSTM that is composed of the forward and backward LSTMs, which can capture previous and future context information. Based on the final forward and backward LSTM outputs, the LSTM-2 is expressed as follows,

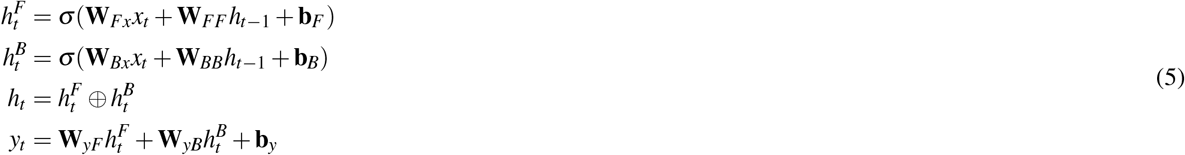

where 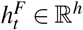 and 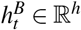 represent the forward and backward outputs. **W** and **b** represent the weight matrix and bias vectors, respectively. The ⊕ denotes the concatenate operating. For the LSTM-2, the units of each layer are 256, and the output of LSTM-2 doubles that of the LSTM-1.

In the ESIDEN, the FC layer plays an important role in effectively learning the nonlinear combination of components extracted from the input features. In the FC module, there are two different FC layers: the FC-1 and the FC-2 layers. As illustrated in Figure 1, the FC-1 layer consists of 256 nodes with ReLU activation operator, that is, *f*(*x*) = *x* if *x* ≥ 0, otherwise *f*(*x*) = 0, and 80% dropout are used in the FC-1 layer when the ESIDEN is trained. While the other FC layer (FC-2) has four outputs with the same Sigmoid activation function 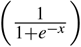 which converts the four outputs of the ESIDEN into real values. Accordingly, the detailed architecture of the ESIDEN is designed based on the above operators, and there are a small number of learnable parameters, about 6.6M in total. Benefiting from those advantages, the ESIDEN developed into a simple and efficient neural network model for predicting the torsioin angles.

### 2.3 Outputs

Both the torsion angles (*ϕ* and *ψ*) are formed by continuously connecting four atoms located in the backbone of the protein, that is, *ϕ* results from atoms *C*_*i*–1_ – *N_i_* – *C_αi_* – *C_i_* while *ψ* is computed from *N_i_* – *C_αi_* – *C_i_* – *N*_*i*+1_ (Figure 1). They are all located in the range [−180°, 180°], and the torsion angles *ϕ* at the N-terminal and *ψ* at the C-terminal are fixed. In our study, the proposed ESIDEN method can simultaneously predict the torsion angles *ϕ* and *ψ* using four outputs (sin (*ϕ*), cos (*ϕ*), sin (*ψ*), and cos (*ψ*)). To remove the effect of angle’s periodicity, we use the sine and cosine values of each torsion angles as targets instead of directly predicting *ϕ* and *ψ*, accordingly, the predicted values of an angle (*P*) is defined as follows,

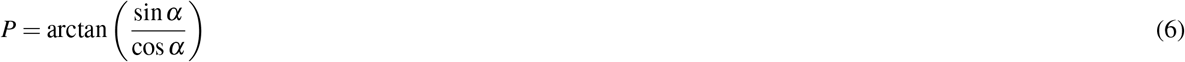

where *α* is a representation of either the angle *ϕ* or *ψ*.

### 2.4 Performance evaluation

We evaluate the accuracy of the predicted torsion angles by the mean absolute error (MAE), which is to measure the average absolute difference between predicted angles (*P*) and experimental values (*E*) over all residues in a protein chain. To reduce the periodicity of an angle and the artificial effect, we take the minimum value between |*P_ij_* – *E_ij_*| and 360° – |*P_ij_* – *E_ij_*|, i.e.

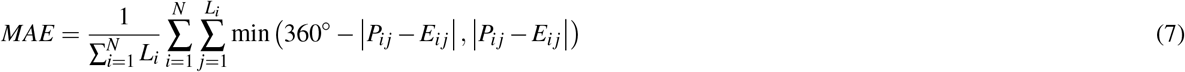

where *N* is the number of protein chains, *L_i_* is the total number of residues in the *i*th protein chain. *P_ij_* and *E_ij_* are the values of predicted and experimental angles of the *j*th residue in the *i*th protein chain, respectively.

## 3 Results

The developed ESIDEN network is implemented in PyTorch v1.7.0^45^, and it is trained on high-performance computational clusters using one NVIDIA GTX2080Ti Graphics Processing Unit (GPU). During the training, we use the Adam optimization algorithm^46^ with a learning rate of 0.001 to optimize the parameters of the ESIDEN network, and the mean square error (MSE) between the predicted and experimental values is defined as a loss function, which used to update the weights and biases of the network. The batch size used in this study is set to 32, and the maximum number of iterations is 5,000.

To demonstrate the performance of the developed ESIDEN, we conduct experiments on two benchmark datasets, independent dataset (D2020) and SPOT-1D dataset^31^ (TEST2016 and TEST2018). On our dataset (D2020, Table S1), we first analyze how much each feature can contribute to improve the prediction accuracy of the ESIDEN and evaluate the importance of each feature. Further, we demonstrate that the four novel features (RE, DC, PSSP, and RBP) can improve the accuracy in predicting the torsion angle when they are combined with the basic features, and the performance of the ESIDEN is accessed by the combinations of different features on the same dataset. On the TEST2016 and TEST2018 datasets (Table S2), we compare the performance of the ESIDEN to those of the best-so-far state-of-the-arts, including Spider3^20^, RaptorX-Angle^26^, SPOT-1D^27^, and OPUS-TASS^8^. We also validate the ESIDEN network on predicting the torsion angles of the fifty-nine CASP TFM targets. On these targets, we compare the predicted torsion angles of the ESIDEN to those of the three best-so-far methods (Spider3, RaptorX-Angle, and SPOT-1D), which are implemented locally using their standalone packages on the same computing resources. Finally, the torsion angles estimated by the ESIDEN are leveraged to demonstrate its ability in predicting the tertiary structures of four representative TFM targets of the fifty-nine proteins in the CASP dataset.

### 3.1 Independent features

In this section, we compare basic features (AA, PP, and PSSM) and novel features (DC, RE, PSSP, and RBP) on the ESIDEN(Figure 1), and we also evaluate the performances measured by the MAE between the predicted and experimental torsion angles on the validation and test dataset of the D2020 (Table S1). As illustrated in Table 1, two of the basic features (AA and PP) achieve comparable MAE of *ϕ* and *ψ* using the developed model, while another basic feature PSSM is better than the two basic features. Compared to the basic features, the MAE *ϕ* and *ψ* of the ESIDEN with the DC is slightly higher than the two basic features (AA and PP), as the DC loses much more information than the two features. Notably, the MAE of the model with either the RE, RBP, or PSSP is lower than those of both the features AA and PP, especially, the features PSSP and RBP achieve comparable MAE to the PSSM, as the PSSP contains a little more noises than the PSSM. Due to the lack of distinguishable characteristics of amino acids, the RBP is slightly worse than that of PSSM and PSSP. Similarly, the predicted accuracy based on the RE is better than that of the DC as DC loses more information than the RE. They all don’t preserve much discernible information about different residues as the PSSM and PSSP, and it could be the reason why PSSM and PSSP provide better predictions by comparing to that of the RE and DC on both torsion angles. Although the RE loses information, the precision of our method based on the RE is still better than that of the two basic features (AA, PP). Accordingly, we find that a single feature is not sufficient to enhance the performance of the proposed method. We further conduct analyses on different combinations of features and measure the prediction accuracy of the ESIDEN on the combined features.

**Table 1.**
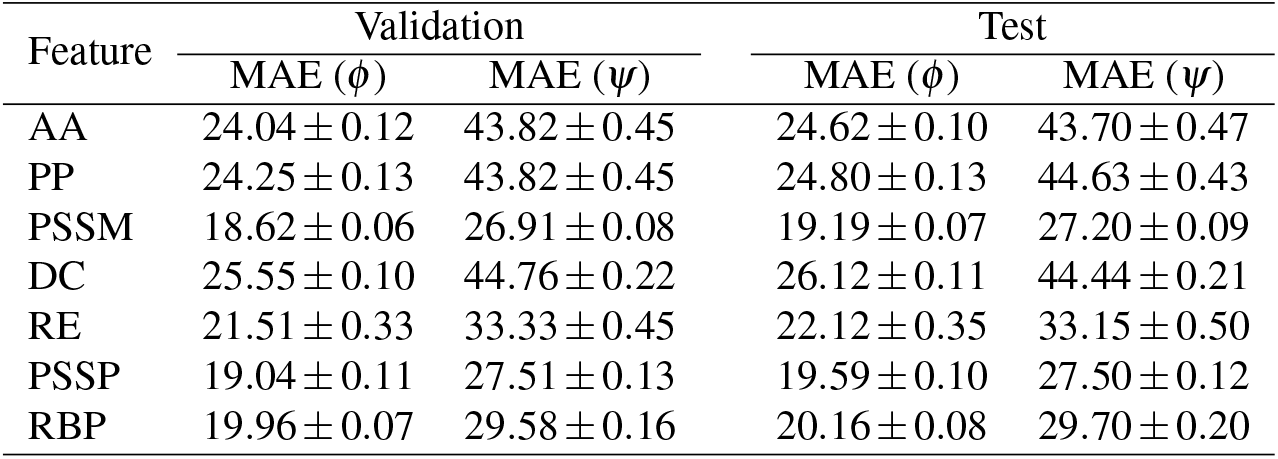
The MAE of each single feature in predicting the torsion angles using ESIDEN on the D2020 dataset

### 3.2 Combined features

To address how joint contribution the novel features make to the prediction, we produce different combinations of all the features, including the basic and new ones (Table 2), to validate the performance of the ESIDEN in predicting the torsion angles (*ϕ* and *ψ*).

**Table 2.** The MAE of combined features in predicting the torsion angles using ESIDEN on the D2020 dataset

| Combined features | Validation |  | Test |  |
| --- | --- | --- | --- | --- |
| | MAE ($\phi$) | MAE ($\psi$) | MAE ($\phi$) | MAE ($\psi$) |
| Basic = PSSM + PP + AA | $17.22 \pm 0.08$ | $25.19 \pm 0.08$ | $18.00 \pm 0.08$ | $25.87 \pm 0.09$ |
| New = RE + DC + PSSP + RBP | $16.64 \pm 0.09$ | $21.52 \pm 0.16$ | $17.24 \pm 0.13$ | $22.10 \pm 0.09$ |
| Basic + DC | $17.05 \pm 0.06$ | $24.88 \pm 0.11$ | $17.76 \pm 0.08$ | $25.33 \pm 0.14$ |
| Basic + RE | $17.18 \pm 0.08$ | $25.06 \pm 0.08$ | $17.82 \pm 0.08$ | $25.38 \pm 0.09$ |
| Basic + PSSP | $16.38 \pm 0.07$ | $23.21 \pm 0.10$ | $16.96 \pm 0.08$ | $23.51 \pm 0.09$ |
| Basic + RBP | $16.32 \pm 0.06$ | $21.08 \pm 0.11$ | $17.02 \pm 0.09$ | $21.70 \pm 0.14$ |
| CbnF = Basic + DC + RE | $17.01 \pm 0.08$ | $24.81 \pm 0.09$ | $17.64 \pm 0.08$ | $25.08 \pm 0.08$ |
| CbnF + PSSP | $16.21 \pm 0.08$ | $22.92 \pm 0.09$ | $16.78 \pm 0.08$ | $23.23 \pm 0.07$ |
| CbnF + RBP | $16.18 \pm 0.09$ | $20.89 \pm 0.08$ | $16.91 \pm 0.09$ | $21.59 \pm 0.09$ |
| CbnF + PSSP + RBP | $15.72 \pm 0.07$ | $20.06 \pm 0.06$ | $15.72 \pm 0.07$ | $19.77 \pm 0.05$ |

As shown in Table 2, we compare the MAE performance of different feature combinations on our validation and test set by using the ESIDEN. Firstly, the combination of all the four novel features (RE + DC + PSSP + RBP) outperforms the basic features (PSSM + PP + AA) with regard to the MAE of the both angles. In particular, the MAE of the angle *ψ* predicted by the novel features is much better than that of the basic features (PSSM, PP, and AA), that is, the MAE (*ψ*) is reduced by about 4 for both the validation and test set. Combined with the basic features, every single new feature can still improve the prediction accuracy on the torsion angles. For example, the RE slightly improves the performance in predicting the angles *ϕ* and *ψ* when compared to that of the combined basic features. The DC outperforms both the basic features and their combination with the RE, and it would be a result of the DC preserves conservation information from protein evolution with much fewer noises as the RE. The combined features (Basic+PSSP) distinctly improve the predicted angle *ϕ*, and it’s worth noting that the combined feature (Basic+RBP) makes a significant contribution to increasing the prediction accuracy of the angle *ψ*. The RE and DC combined with the basic features (Basic+DC+RE) are a little better than that of each in fusion with the basic features. Similar performances are achieved by the ESIDEN based on the PSSP and RBP combined with the basic features, interestingly, the RBP in the fusion of the basic features plays a significant role in improving the prediction accuracy of the angle *ψ*. Notably, the fusion of all the basic and the new features remarkably decreases the MAEs from 18.00 to 15.72 and 25.87 to 19.77 for the angles *ϕ* and *ψ*, respectively, by comparing to those of the basic features (Figure 2). These promising results demonstrate that the four novel features can significantly improve the prediction accuracy of the torsion angles, and they can make the prediction much better when combined with the basic features. Therefore, we use the combination of basic and novel features as the input features to build a small but efficient network model in this study.

**Figure 2.**
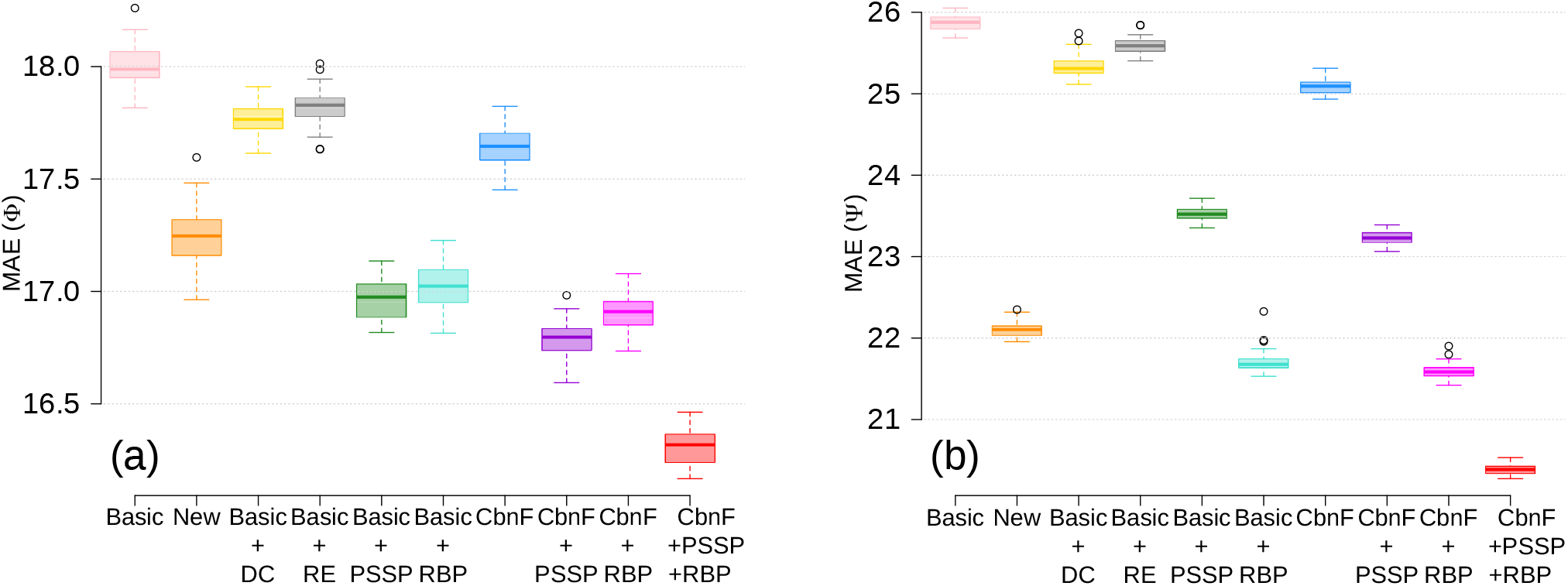
Comparison of prediction performance using different combinations of features on the D2020 dataset. The MAE as measurements between the predicted and experimental values of (a) the torsion angle *ϕ* and (b) the torsion angle *ψ*, respectively.

### 3.3 Comparisons to other the-state-of-the-arts

In this section, we compare the MAE performance of the proposed ESIDEN to those of other the best-so-far methods (Spider3^20^, RaptorX-Angle^26^, SPOT-1D^27^ and OPUS-TASS^8^) on the widely used datasets, TEST2016 and TEST2018.

As shown in Table 3, on the TEST2016 dataset, the Spider3 and the RaptorX-Angle achieve comparable MAE of the angle *ψ*, but the Spider3 slightly outperforms the RaptorX-Angle in terms of *ϕ* MAE. The hybrid model, OPUS-TASS, performs better measured by the MAE of *ϕ* than the SPOT-1D, although the MAE of the predicted *ψ* by the SPOT-1D is a little better than that of the OPUS-TASS. Our method, the ESIDEN, achieves the best MAE accuracy on the TEST2016 dataset, as shown, it obtains better MAE of the angle *ϕ* than that of the OPUS-TASS on the TEST2016 dataset, and the MAE of the angle *ψ* inferred by the ESIDEN is much better than all the other four compared methods by reducing the MAE more than 5 degrees. On the TEST2018 dataset, the RaptorX-Angle has underperformed the other compared methods with regard to the MAE of both *ϕ* and *ψ*. The SPOT-1D and OPUS-TASS obtain similar MAEs of the angles *ϕ* and *ψ*, and both of them are better than that of the Spider3. On the same dataset, the ESIDEN achieves a better MAE of *ϕ* than those of all the other four compared methods. It’s worth noting that the MAE of the angle *ψ* is much decreased by the ESIDEN compared to those of the rest methods. The results demonstrate that the ESIDEN outperform among the compared methods, especially in terms of the *ψ* MAE, and it significantly improves the prediction accuracy of the angle *ψ*, decreasing the MAE by 3.8 degrees over the OPUS-TASS (the second best) on both of the benchmark datasets. Benefiting from evolutionary signatures, the delicately designed architecture of the ESIDEN accounts for the distinguishing outperformance on accurately predicting the torsion angles.

**Table 3.** Performance of different methods on the TEST2016 and TEST2018.

| Method | Test2016 |  | Test2018 |  |
| --- | --- | --- | --- | --- |
| | MAE ($\phi$) | MAE ($\psi$) | MAE ($\phi$) | MAE ($\psi$) |
| Spider3* | 17.88 | 26.66 | 18.38 | 28.10 |
| RaptorX-Angle* | 18.08 | 26.68 | 21.01 | 35.95 |
| SPOT-1D* | 16.27 | 23.26 | 16.89 | 24.87 |
| OPUS-TASS<sup>†</sup> | 15.78 | 24.46 | 16.40 | 24.06 |
| ESIDEN | 15.48 | 19.25 | 16.00 | 20.28 |

\*The results are obtained from the SPOT-1D<sup>27</sup> paper
<sup>†</sup>The results are collected from the OPUS-TASS<sup>8</sup> paper

### 3.4 Validations on the CASPs datasets

The CASP datasets are widely used to evaluate the performance of different predictors on structural informatics. Here, we collect 59 TFM targets from the recent CASPs (CASP11, CASP12, CASP13, and CASP14, Table S3), and the proposed ESIDEN is validated by comparison to the best-so-far methods (Spider3^20^, RaptorX-Angle^26^ and SPOT-1D^27^) in predicting the torsion angles. As shown in Table 4, on the four CASP datasets, the the MAE performance of ESIDEN is slightly better than those of all the compared methods, while the decrement of its MAE of *ψ* more than 2 degrees, except 0.5 degrees on the CASP13, by comparing to those of the others. On the CASP11, CASP12, CASP13 and CASP14 datasets, the performance of our method consistently outperforms the performances of all the other three methods, especially, the *ψ* MAE of the ESIDEN is distinguishably better than those of the other methods. The average torsion angle prediction errors (measured by the MAE) of the ESIDEN are demonstrated on the fifty-nine TFM targets across eight types of secondary structures (Q8) (Figure 3). There are no secondary structures (5-turn helix I, and bend S) in the CASP datasets, and the result shows that the prediction errors of both *ϕ* and *ψ* for H (helix) are the lowest by comparing other Q8 types, yet the secondary structures B/T/C are higher, as variational coiled coils and single *β* strands are not stable as the *α*-helix.

**Figure 3.**
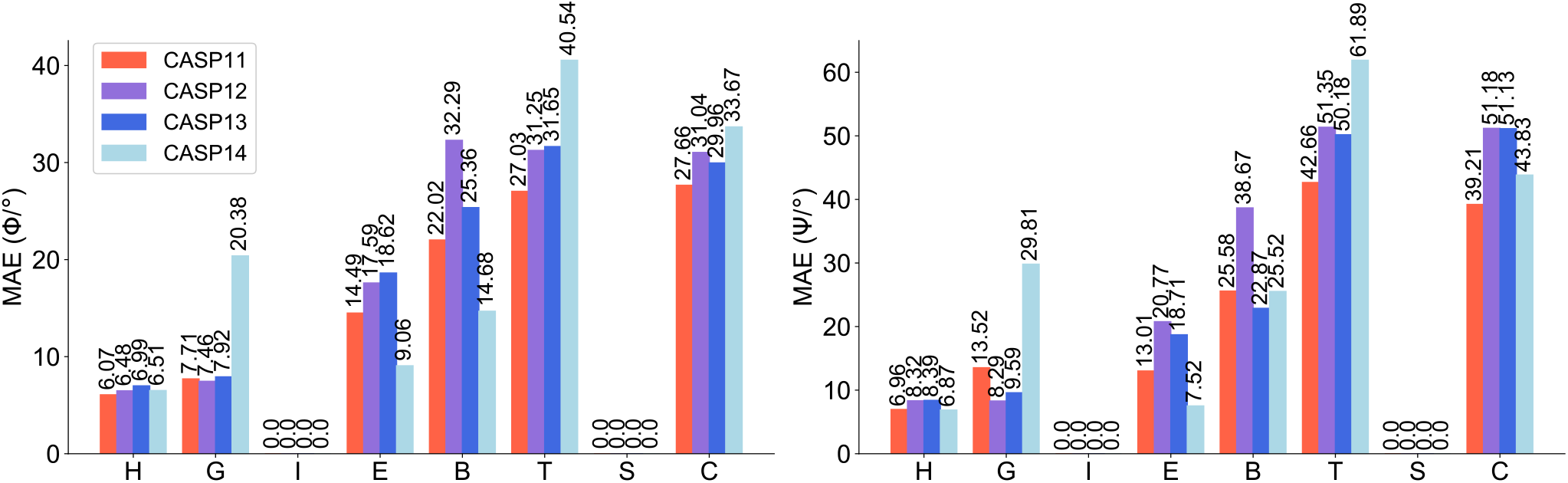
The MAE of the torsion angles (*ϕ* and *ψ*) on the 8-class secondary structures for the TFM targets of the CASPs.

**Table 4.** Comparison among different methods on the TFM targets of the CASP11, CASP12, CASP13, and CASP14

| Method | CASP11 (27) |  | CASP12 (11) |  | CASP13 (13) |  | CASP14 (8) |  |
| --- | --- | --- | --- | --- | --- | --- | --- | --- |
| | MAE ($\phi$) | MAE ($\psi$) | MAE ($\phi$) | MAE ($\psi$) | MAE ($\phi$) | MAE ($\psi$) | MAE ($\phi$) | MAE ($\psi$) |
| Spider3* | 19.19 | 34.63 | 21.14 | 34.92 | 22.48 | 38.46 | 23.41 | 38.79 |
| RaptorX-Angle* | 20.33 | 40.05 | 21.71 | 38.22 | 22.73 | 41.18 | 24.95 | 48.06 |
| SPOT-1D* | 18.54 | 25.77 | 20.21 | 31.71 | 22.60 | 34.28 | 23.42 | 33.44 |
| ESIDEN | 17.25 | 23.30 | 19.94 | 28.86 | 22.15 | 32.40 | 23.01 | 29.96 |

\*The results are obtained locally using the Spider3, RaptorX-Angle, and SPOT-1D standalone packages, respectively

We model four representative TFM targets of the 59 TFM targets in the CASPs using the predicted torsion angles (Figure S2–S5), including T0968s2-D1, T0986s1-D1, T0957s2-D1, and T0969-D1 (Table S4), and the inferred angles are used as a constraint to launch the folding module in the *Leri* software^38^. Ten folding simulations are launched for each target. For the targets T0986s1-D1, T0968s2-D1, and T0957s2-D1, we implement ten folding simulations with 100,000 iterations and obtain 200 structures with lower energy from each trajectory. As the number of residues of the target T0969-D1 is large, we conduct 200,000 iterations and obtain 200 structures with lower energy from each trajectory. The largest cluster is obtained over the two thousand structures of each target, and the centroid structure in the cluster is compared to its native structure, as well as the best-so-far structure (Figure 4). The root-mean-square deviation (RMSD) and TM-score of each target are computed by the TMscore software^47^, and they are presented in Table S4. Although the coiled structures of either the best-so-far or the centroid structures are not always in agreement with those in the native structures, the secondary structures (*α*-helix and *β*-strand) in both the best-so-far and centroid structures share the same ones as those in the native structure of each target. Accordingly, the predicted torsion angles can accelerate protein folding and structure prediction, and they can also improve the structural precision.

**Figure 4.**
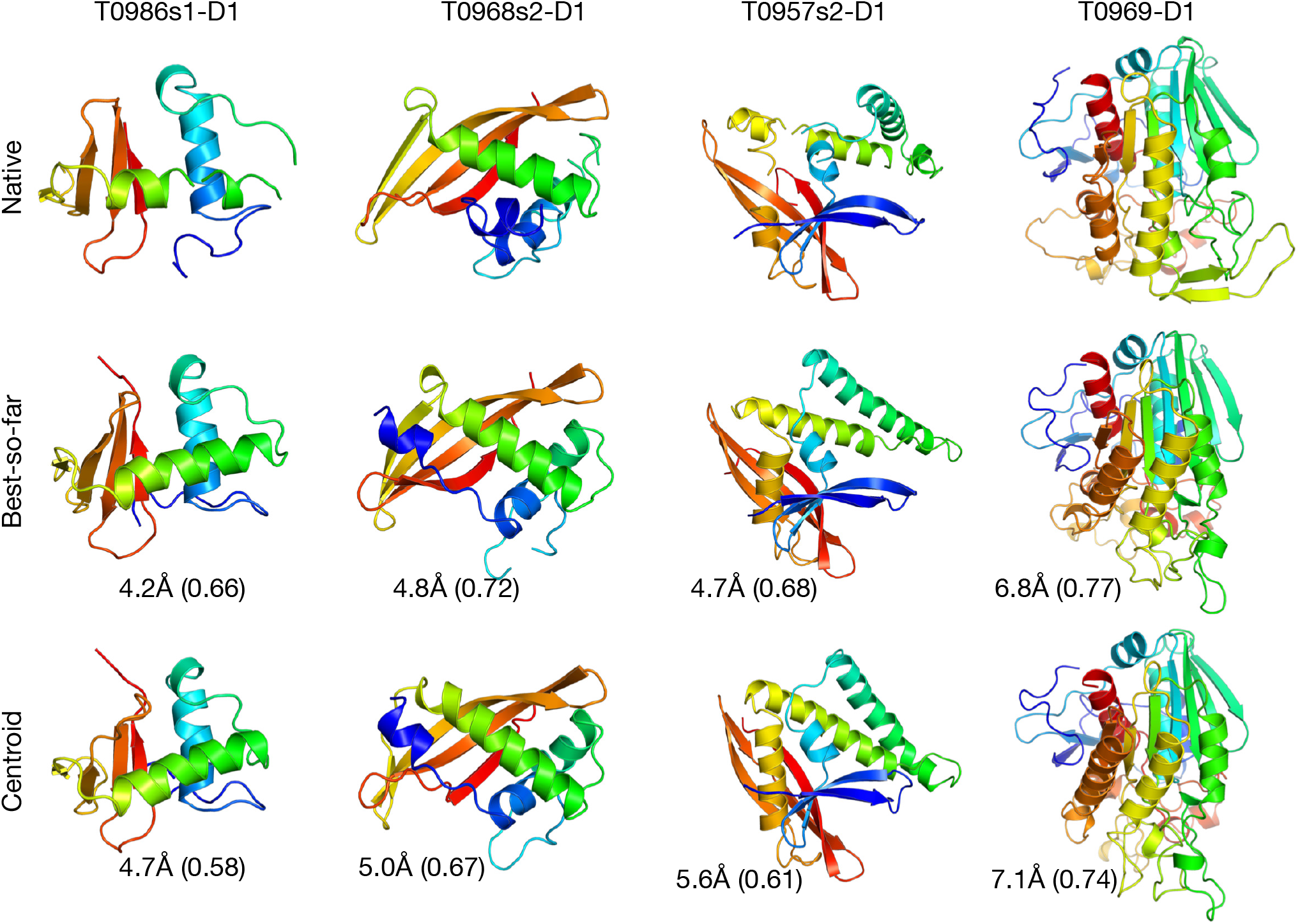
The predicted structures of four representative TFM targets using the torsion angles estimated by the ESIDEN.

## 4 Discussion

The classical features extracted from a protein sequence are powerful input data for the ML-based methods, which predict protein structural properties, such as residue contacts, residue distance maps, backbone torsion angles, solvent accessible surface area, protein-protein interaction, and protein function in computational biology. They have several advantages for characterizing amino acid types and sequential order of amino acids that can be able to specify structural properties. Therefore, accurate prediction of structural properties can provide valuable information to infer protein tertiary structure and function. Recently, incorporation of pre-estimated structural features, e.g., secondary structure and solvent accessibility, into the ML-based methods has been a practical way to improve the predictions for either protein structure or function. Nevertheless, it still remains challenging to further improve the prediction accuracy of these key structural properties, such as the torsion angles, merely based on the classical features. Extracting efficient features from the large-scale sequence profiles increasingly advance the capabilities of computational methods, especially DL-based algorithms, to draw out knowledge and address important biological questions, e.g., the quantitative relationship between protein structure and its function.

In this study, we propose an evolutionary signatures-driven deep neural network-based system (ESIDEN) and four novel features computed from evolutionary signatures and the Ramachandran basins for predicting protein torsion angles. The ESIDEN adopts a straightforward architecture of a small number of learnable parameters, and yet it achieves high accuracy in the torsion angle prediction in comparison to other state-of-the-art approaches. Furthermore, we also developed four novel features that are derived from protein evolutionary signatures to improve the performance of ESIDEN. Generally, it is more difficult to predict the angle *ψ* than *ϕ*, as the diversity of *ψ* results from widely different secondary structures for variational amino acids in proteins. In contrast to existing methods that leverage the classical features by combining other features to substantially improve performance, the ESIDEN is built upon the different features, especially, the ESIDEN with only the RBP can still achieve similar performance by comparison to that of PSSM. The results demonstrate that the newly developed features distinguishably improve the accuracy in predicting the angle *ψ*. Moreover, the recurrent architecture in the ESIDEN, can capture sequential motifs hidden in the amino acid residues and their neighbors. On the TEST2016, TEST2018, and the CASP datasets, we demonstrate the ESIDEN achieves higher precision of predictions on the torsion angles by comparison to the best-so-far methods. The accurately predicting the torsion angles is the result of the efficient architecture of the ESIDEN and the new features that contribute to the classical ones.

Limitations of the model include biases that arise from features filtered by the dimension-reduction methods on the Ramachandran basins, which may result in computationally intractable reduction if improper methods are used and evolutionarily younger families of limited diversity that result in many noises to the feature PSSP. Although incorporating evolutionary signatures into the ESIDEN results in a practical improvement over other methods, challenges remain in the precise interpretation of the model.

The success of the ESIDEN is based on deep learning at recapitulating large-scale data from protein sequence profiles. For example, the Ramachandran basins could robustify other deep learning-based methods for many applications in predicting protein contacts/distances, secondary/tertiary structure, and designing proteins. We anticipate that the ESIDEN and the new features developed in this study can be utilized by other deep-learning-based methods applications ranging from drug discovery to protein design. The consistency of our estimated torsion angles with the authentic angles highlights how the inclusion of evolutionary signatures will facilitate more accurate inferences and aid the prediction/determination of the tertiary structures of protein sequences.

## Acknowledgements

We acknowledge Dr. Arun T. John Peter for discussion and proofreading the manuscript. We also thank all members in the Ding’s Group for helpful discussions. XYC, TJSG, and NJC are supported by the Leri Ltd., UK.

## Competing interests

Potential conflicts of interest. NJC (YZ) is a founder of Leri Ltd based in Oxford, UK. All other authors report no conflicts of interest relevant to this article.

## Data and code availability

The extracted features and trained models can be accessed and freely downloaded at https://kornmann.bioch.ox.ac.uk/leri/resources/download.html.

